# Dihydroartemisinin broke immune evasion through YAP1/JAK1/STAT1, 3 pathways to enhance anti-PD-1 therapy in hepatocellular carcinoma

**DOI:** 10.1101/2021.11.30.470572

**Authors:** Qing Peng, Shenghao Li, Xinli Shi, Yinglin Guo, Liyuan Hao, Zhiqin Zhang, Jingmin Ji, Yanmeng Zhao, Caige Li, Yu Xue, Yiwei Liu

## Abstract

The efficacy of anti-PD-1 therapy is not as expected in patients with hepatocellular carcinoma (HCC). Yes-associated protein 1 (YAP1) was overexpressed and activated in HCC. This study aimed to investigate the potential mechanism and inhibitor of YAP1 on immune evasion, and promote anti-PD-1 therapy in HCC. Here, we showed that dihydroartemisinin (DHA), an FDA approved drug, directly suppressed YAP1 expression, leading to break immune evasion in liver tumor niche, characterized by decreased PD-L1 in liver tumor cells and increased CD8^+^ T cell infiltration. Mechanismly, YAP1 is not only directly related to PD-L1, but also involved in activating the JAK1/STAT1, 3 pathways. Moreover, *Yap1* knockout elevated CD4^+^ and CD8^+^ T cells in liver tumor niche of *Yap1*^LKO^ mice. Consistently, verteporfin, YAP1 inhibitor, decreased TGF-β in liver tumor niche and exhausted CD8^+^ T cells in spleen. Furthermore, DHA combined with anti-PD-1 treatment promoted CD4^+^ T cell infiltration in the spleen and CD8^+^ T cells in tumor tissues. Thus, we provide a new combined therapeutic strategy for anti-PD-1 with DHA, a potent YAP1 inhibitor, in HCC.

## Introduction

Hepatocellular carcinoma (HCC) is the fourth most common cause of cancer-related death in the world (Lei et al. 2019). Immune checkpoint inhibitor (ICI) therapy, particularly antibodies targeting the programmed cell death-1 (PD-1)/programmed cell death ligand-1(PD-L1) pathway, has shed light on the survival of HCC patients. However, the objective response rate is only ~20% during PD-1/PD-L1 blockade therapy in cancers (Xu-Monette et al. 2017). In the tumor microenvironment, PD-L1 on tumor cells is a key transmembrane molecule that governs the crosstalk with tumor-infiltrating CD8^+^ cytotoxic T lymphocytes (CTLs), which played an important role in the ICI therapy.

PD-L1 is highly expressed in HCC tissues compared to adjacent tissues, which positively correlated with poor prognosis and invasion (Calderaro et al. 2016). PD-L1 on liver tumor cells is induced by interferon gamma (IFN-γ) secreted from CD8^+^ CTLs in tumor microenvironment, drives T cell exhaustion, and forms a negative feedback loop, leading to immune evasion (Huang et al. 2017). Therefore, there is an urgent need to develop the mechanism negatively regulates PD-L1 expression in HCC.

Yes-associated protein 1 (YAP1), a key effector in Hippo pathway, directly binds to PD-L1 promoter and promotes PD-L1 transcription in lung cancer PC9 cells (Lee et al. 2017). Overexpression and nuclear localization of YAP1 is about 50% in HCC clinical specimens (Li, Li, and Zhou 2017). YAP1 overexpression recruits inhibitory immunocyte including tumor associated macrophages (tumor-associated macrophages (TAMs), M2 type) (Guo et al. 2017), myeloid-derived suppressor cells (MDSCs) (Wang et al. 2016) and Tregs (Fan et al. 2017). In addition, the phosphorylation STAT1 (T727) or STAT3 (Y705) also bound to the PD-L1 promoter and induced PD-L1 expression (Sasidharan Nair et al. 2018). However, the relationship between YAP1 and STAT1, 3 in regulating PD-L1 of HCC remains unclear.

Dihydroartemisinin (DHA), approved by FDA as an anti-malarial drug, is a derivative of artemisinin extracted from *artemisia annua*. In addition, DHA inhibited the expression of PD-L1 by inhibiting TGF-β, STAT3 and PI3K/AKT signaling pathways in non-small cell lung cancer (Zhang et al. 2020). Our previous study showed that DHA inhibited cell proliferation in human hepatocellular carcinoma HepG2215 cells (Shi et al. 2019) and promoted p-STAT3 (Y705) nuclear localization in human tongue squamous cell carcinoma Cal-27 cells (Shi et al. 2017). However, the relationship between DHA and YAP1 in the immune microenvironment is unknown in HCC.

Here we showed that the anti-PD-1 treatment increased YAP1 expression in liver tumor cells and the exhausted CD4^+^ and CD8^+^ T cells in blood and spleen. YAP1 knockdown/knockout decreased PD-L1 expression and promoted CD8^+^ T cells infiltration in liver tumor niche. Mechanistically, YAP1 prompted PD-L1 expression by JAK1/STAT1, 3 pathways in liver tumor cells. Interestingly, DHA acted as YAP1 inhibitor and enhanced the effect of anti-PD-1 therapy in HCC.

## Results

### YAP1 expressed differently in tumor and para-tumor tissues from HCC patients

We investigated the expression patterns of YAP1 in different tumors by TIMER database. *YAP1* was upregulated in CHOL, COAD, GBM, LIHC and STAD (Fig. 1A). However, *YAP1* was downregulated in BLCA, BRCA, KICH, KIRC, KIRP, LUAD, LUSC, PCPG, PRAD and UCEC (Fig. 1A). The result showed that YAP1 had different expression patterns in different tumors. Next, the TCGA database showed that *YAP1* was also upregulated and constantly increased in HCC tumor tissues of different stages compared with normal tissues (Fig. 1B). In addition, the expressions of YAP1 in tumor and para-tumor tissues were analyzed by the clinical tissue microarray. Representative pictures of YAP1 expression ranged from negative to moderately positive (Fig. 1C). The data showed that YAP1 (Supplementary Table 2) had no correlation with sex, age, histological grade, maximum diameter of tumor (cm), intrahepatic satellite focus, lymphatic metastasis, extrahepatic metastasis, virus infection, HBV and cirrhosis *(P*>0.05). Notably, some study showed the survival of tumor cells depended on the relative activity of YAP1 in tumor cells and their surrounding tissues (Moya et al. 2019). We found that the ratio of YAP1 score (para-tumor/tumor tissues) was significantly negatively correlated with the expression of YAP1 in tumor tissues (R=-0.64, *P*=0.001) (Fig. 1D, Fig. S1). Thus, different expressions of YAP1 were in HCC tumor and para-tumor tissues.

**Figure 1.**
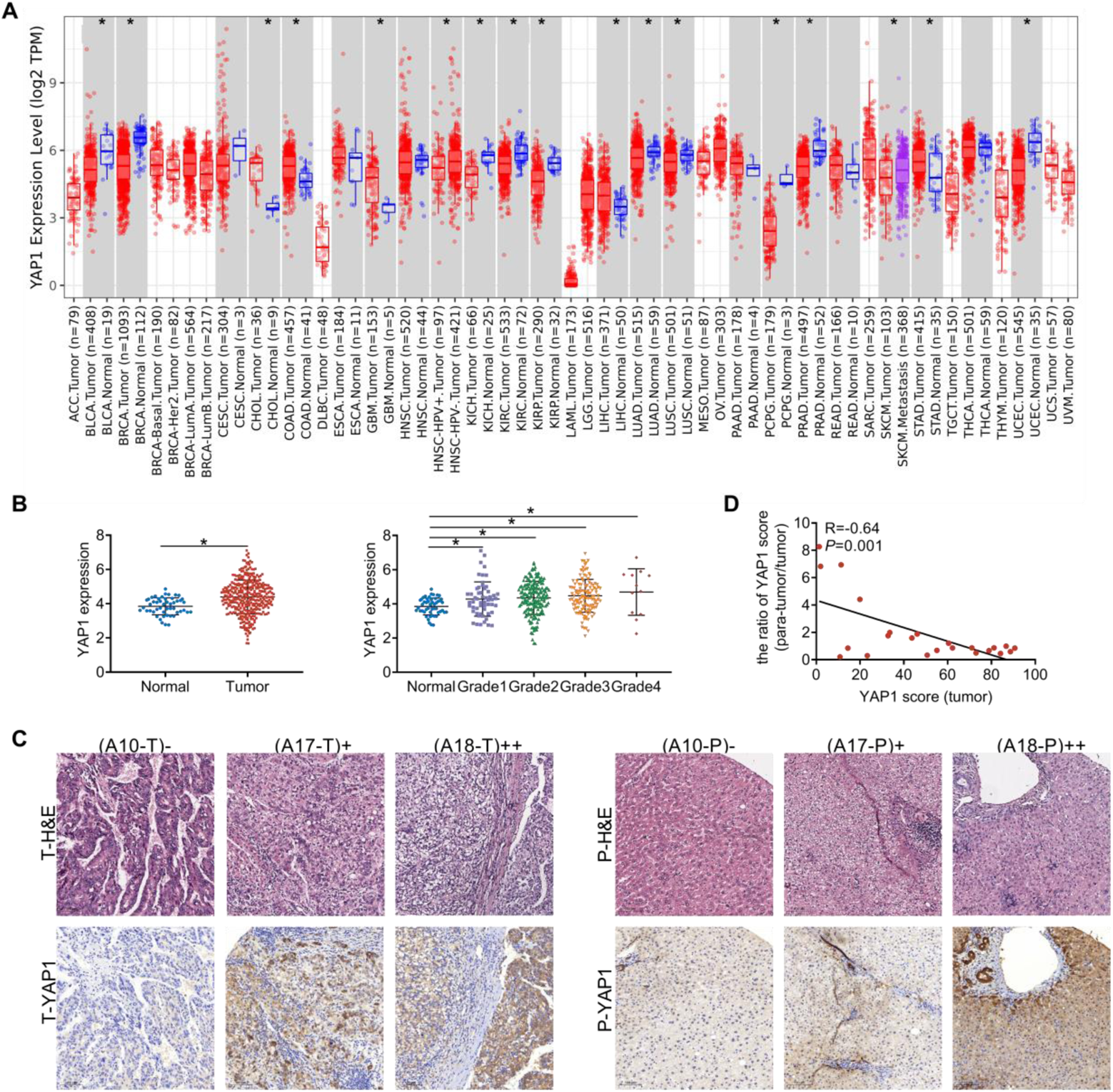
Different expression of YAP1 in tumor and para-tumor tissues from HCC patients. A. Pan-cancer analysis of *YAP1* mRNA expression levels in different tumors by TIMER database. \**P*< 0.05 vs corresponding normal control group. B. The mRNA expression levels of *YAP1* in HCC tumor (T, n=371) and normal tissues (N, n=50), and *YAP1* expression in different HCC grades based on the TCGA database. \**P*< 0.05 *vs* the normal tissues. C. YAP1 expression in tumor (T) and para-tumor tissues (P) by the HCC tissue microarray. A 0, A17 and A18 indicate patient number. D. Analysis the relevance of the ratio of YAP1 (para-tumor /tumor tissues) with clinical tissue microarray.

### DHA inhibited YAP1 expression in liver tumor cells of mice

Our previous study showed DHA inhibited cell growth in HepG2(Hao et al. 2021) and HepG2215 cells (Shi et al. 2019). To further study the effect of DHA on liver tumor *in vivo*, C57BL/6 mice with liver tumors *in situ* were induced by DEN/TCPOBOP (Li et al. 2020, Bergmann et al. 2017). We observed that DHA reduced tumor volume and liver index, but had no significant effect on serum ALT and body weight (Fig. 2A). These results suggested that DHA inhibit liver tumor growth *in vivo*. Moreover, DHA reduced YAP1 expression in the tumor tissues, consistent with that of verteporfin group (Fig.2B). The result suggested that DHA suppressed YAP1 expression in liver tumor. We also showed that anti-PD-1 decreased the tumor volume, liver index and ALT in serum (Fig. 2A). However, we found that anti-PD-1 increased YAP1 expression in tumor and para-tumor tissues compared with DMSO (Fig. 2B). These results supported that anti-PD-1 promoted YAP1 expression in tumor cells. Notably, tumor volume was reduced in DHA combined with anti-PD-1 group (mark as DHA +anti-PD-1) compared with DHA or anti-PD-1 group alone (Fig. 2A). The result suggested that DHA promoted anti-PD-1 treatment effect *in vivo*. Interestingly, YAP1 expression decreased in tumor, while increased para-tumor tissues compared with DMSO (Fig. 2B).

**Figure 2.**
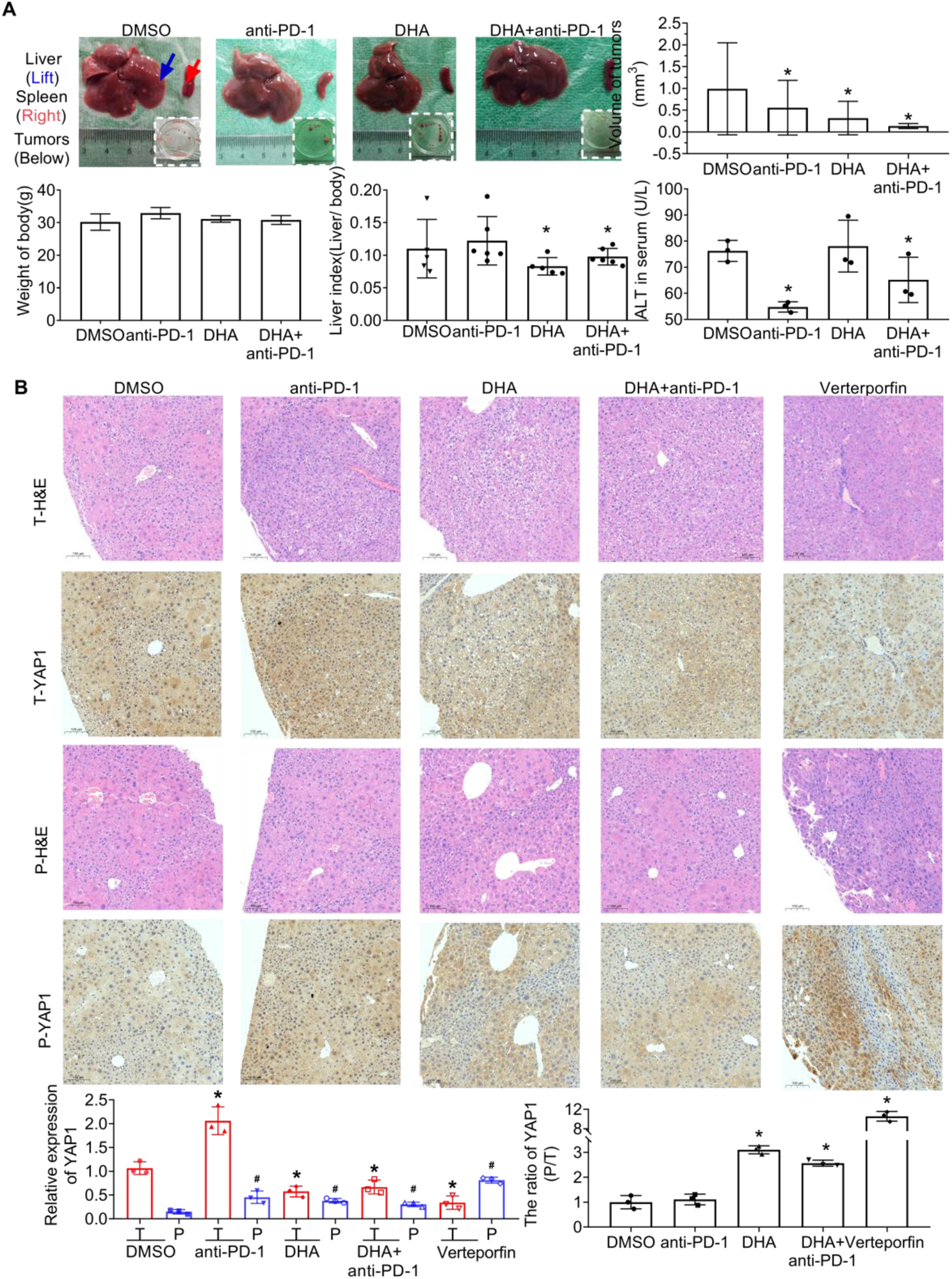
DHA inhibited YAP1 expression in liver tumor mice. A. Representative images of the liver tumor, and tumorigenesis of DHA (25mg/kg) and/or anti-PD-1 (10mg/kg) in DEN/TCPOBOP-induced liver tumor C57BL/6 mice (n=6). \**P* <0.05 *vs* DMSO group. B. IHC staining results of YAP1 in liver tumor (T) and para-tumor (P) in mice with liver tumors *in situ*. \**P* <0.05 *vs* liver tumor tissues, and #*P* <0.05 *vs* para-tumor tissues from DMSO group.

Tumor growth depends on the relative activity of YAP1 in tumor and para-tumor tissues (Moya et al. 2019). We observed that the ratio of YAP1 expression (para-tumor /tumor tissues) was also increased in DHA, DHA+anti-PD-1, and verteporfin groups compared with DMSO (Fig. 2B). These data suggested that the ratio of YAP1 expression can better represent the effect of DHA treatment on tumor growth than YAP1 expression in tumors.

### *Yap1* knockout inhibited liver tumor growth *in vivo*

*Yap1*^LKO^ mice were knockout exon 3 of the *Yap1* gene in liver cells by CRISPR/Cas9 (Fig. 3A). Then, DEN/TCPOBOP was used to induce liver tumor in *Yap1*^flox/flox^ and *Yap1*^LKO^ mice (Bergmann et al. 2017, Li et al. 2012). *YAP1* knockout decreased the maximal tumor size and the numbers of macroscopic tumors in *Yap1*^LKO^ mice (Fig. 3B). Meanwhile, liver index decreased, spleen index increased, but kidney index did not change significantly in *Yap1*^LKO^ mice (Fig. 3B).

**Figure 3.**
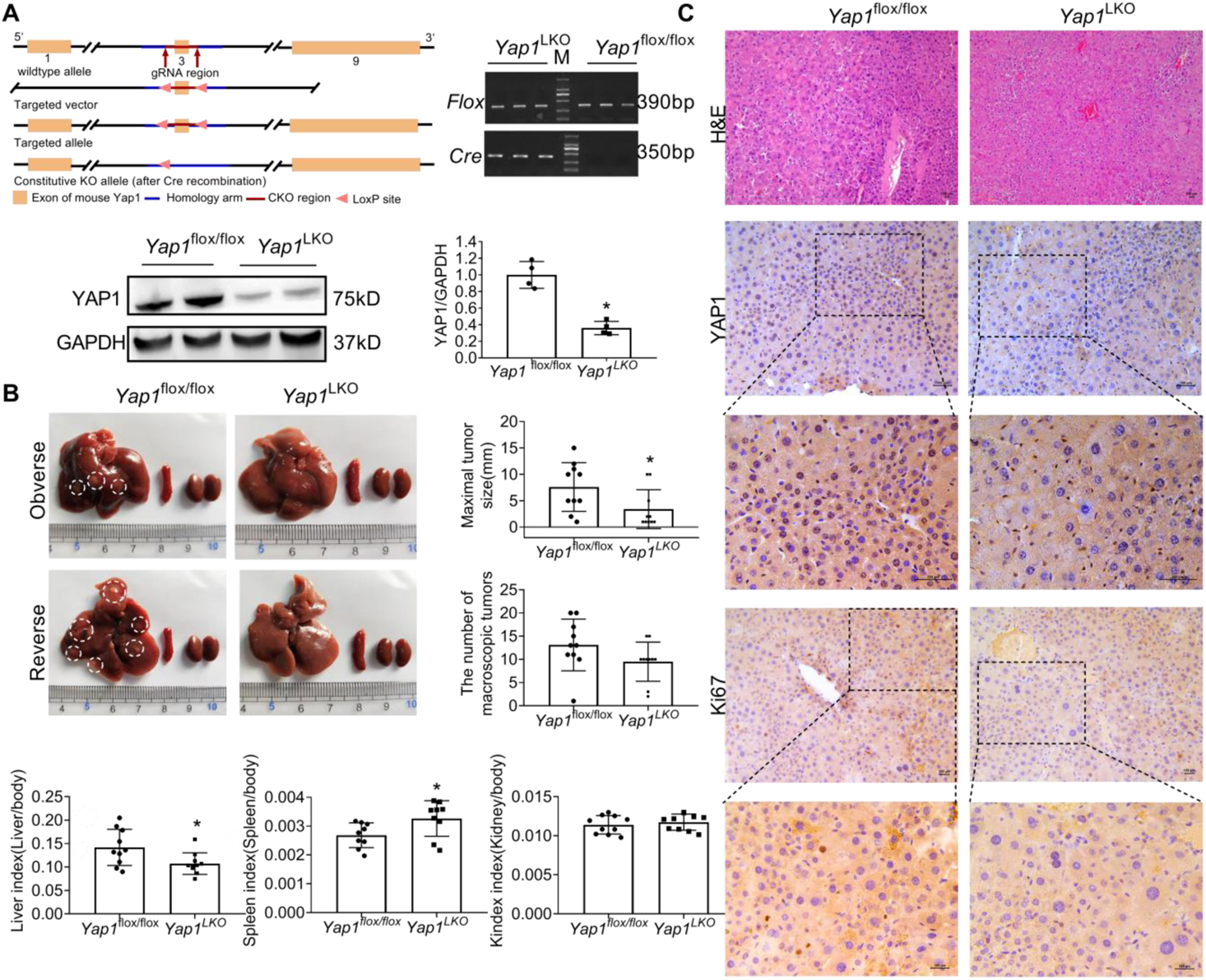
*Yap1* knockout inhibited liver tumor growth *in vivo*. A. Diagram of *Yap1* knockout, genotype identification from the tail, and YAP1 expression of the liver in *Yap1*^LKO^ mice. B. Effect of *Yap1* on tumorigenesis of liver during DEN/TCPOBOP induced tumor in mice (n=9-12). E. IHC staining results of YAP1 and Ki67 expression of liver tumor in *Yap1*^LKO^ mice. \**P* < 0.05.

Consistently, YAP1 was mainly localized in nucleus of liver tumor cells in *Yap1* ^flox/flox^ mice (Fig. 3C). Furthermore, Ki-67 reduced in *Yap1*^LKO^ mice (Fig. 3C). These results indicated that *YAP1* knockout inhibited liver tumor growth *in vivo*.

### DHA directly inhibited liver tumor growth through YAP1 in *Yap1*^LKO^ mice

An increase in reactive oxygen species (ROS) may also contribute to YAP1 activation in human HCC cells (Cho et al. 2020). Our previous study showed that DHA promoted oxygen species (ROS) production in HepG2215 cells (Shi et al. 2019).Further, to verify whether DHA directly inhibited liver tumor growth through YAP1, we treated with DHA in *Yap1*^LKO^ and *Yap1*^flox/flox^ mice with liver tumors. We found that the numbers and maximal size of tumors (Fig. 4A) were reduced in DHA group (2.5±1.7mm) compared with DMSO group (6.0±3.6mm), and Ki67 expression was decreased (Fig. 4B) in *Yap1*^flox/flox^ mice. The results showed that DHA inhibited the tumor growth.

**Figure 4.**
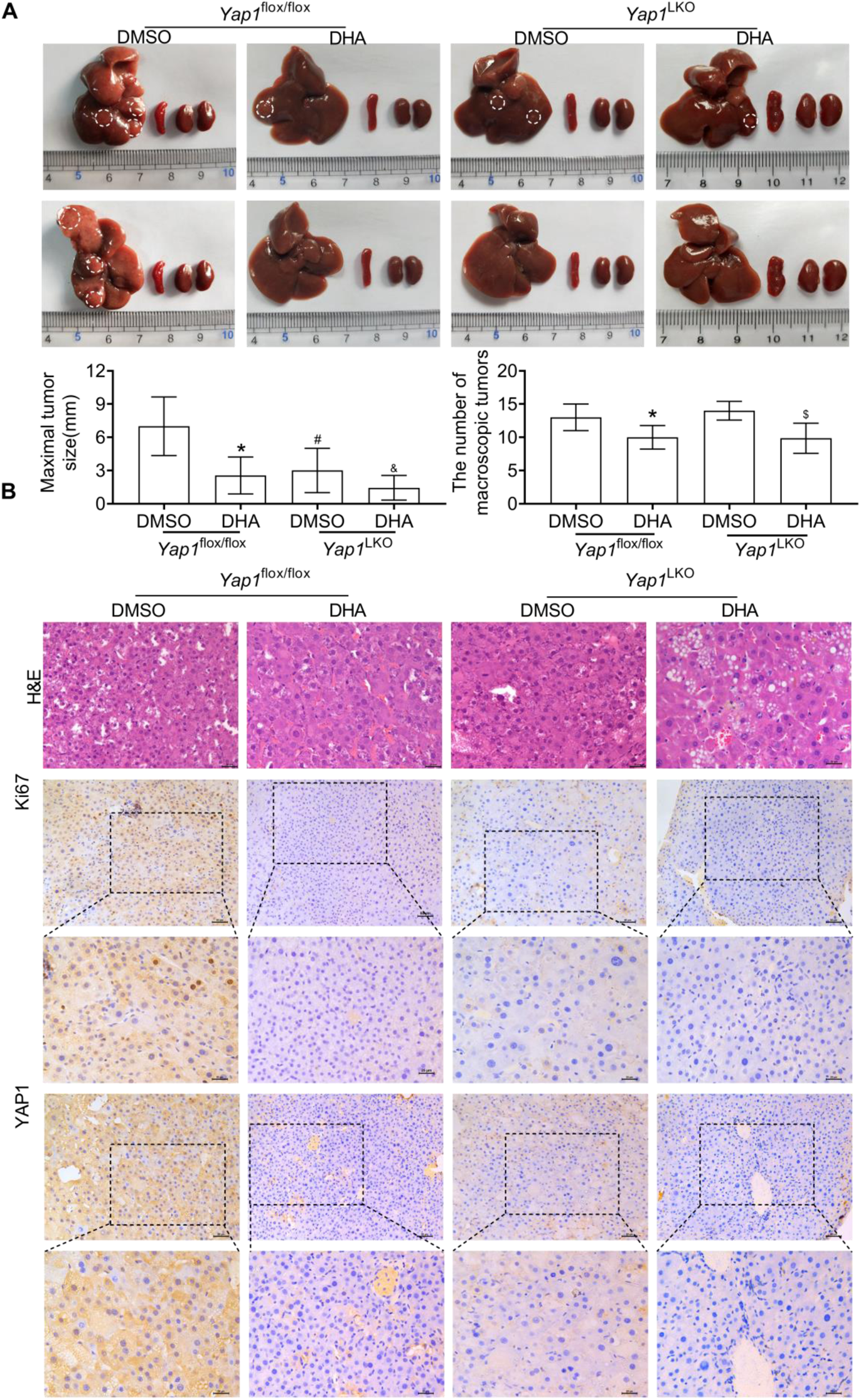
DHA directly inhibited liver tumor growth through YAP1 in *Yap1*^LKO^ mice. A. Effect of DHA on tumorigenesis of liver tumor in *Yap1*^LKO^ mice (n=6). \**P* < 0.05 *Yap1*^flox/flox^+DMSO *VS* Yap1 ^flox/flox^ +DHA; ^#^*P* < 0.05 *Yap1*^flox/flox^ +DMSO *VS* Yap1^LKO^+DMSO;^&^*P* < 0.05 *Yap1*^flox/flox^ +DMSO *VS* Yap1^LKO^+DHA; ^$^*P* < 0.05 *Yap1*^LKO^ +DMSO *VS* Yap1^LKO^+DHA; B. IHC staining of YAP1 and Ki67 in liver tumors of *Yap1*^LKO^ mice.

However, the maximal size (Fig. 4A) and Ki67 expression (Fig. 4B) showed no significant difference in DHA group compared with DMSO group in *Yap1*^LKO^ mice, only tumor numbers decreased (Fig. 4A). Moreover, DHA treatment did not reduce the numbers and maximal size of tumors (Fig. 4A) in *Yap1*^LKO^ mice compared with *Yap1*^flox/flox^ mice, and little difference in Ki67 expression (Fig. 4B) between the two groups. These results suggested that DHA directly inhibited tumor growth through YAP1.

DHA treatment significantly reduced YAP1 expression in *Yap1*^flox/flox^ mice compared with DMSO (Fig. 4B). We detected that YAP1 expression was no significant difference between DHA and DMSO treated *Yap1*^LKO^ mice (Fig. 4B). These results showed that DHA did not restore YAP1 expression.

### YAP1 promoted PD-L1 to immune evasion by JAK1/STAT1, 3 pathways

YAP1 directly bind to the promoter of PD-L1 (Kim et al. 2018). To investigate the effect of YAP1 on PD-L1, we knocked down *YAP1* by CRISPR/Cas9 in HepG2215 cells (Fig. 5A) (Li et al. 2020). The downstream genes cellular communication network factor 1 (*CYR61*) and cellular communication network factor 2 (*CTGF*) were decreased in sh*YAP1*-HepG2215 cells (Fig. S2A). Interestingly, the expression of WW domain containing transcription regulator 1 (TAZ), transcriptional coactivator of YAP1 in Hippo pathway (Yu, Zhao, and Guan 2015), was not significantly altered in sh*YAP1*-HepG2215 cells (Fig. S2B). The result showed that YAP1 knockdown did not affect TAZ expression.

**Figure 5.**
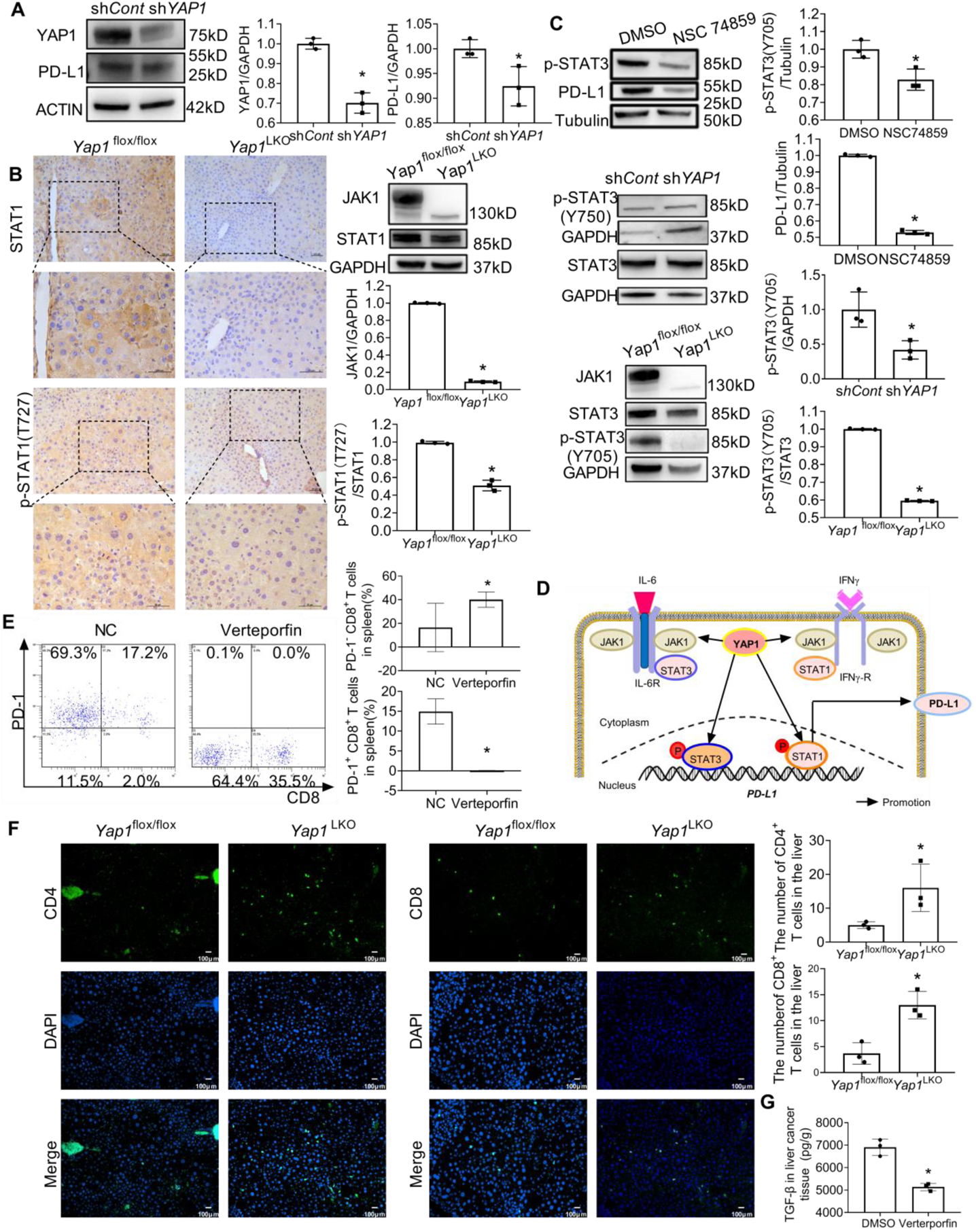
YAP1 promoted PD-L1 expression to immune evasion by JAK1/STAT1, 3 pathways. A YAP1 and PD-L1 expressions in sh*YAP1*-HepG2215 cells.\**P* < 0.05. B. IHC staining and Western blot results of JAK1, STAT1 and p-STAT1 (T727) in liver tumor of *Yap1*^LKO^ mice. \**P* < 0.05. C. Western blot results of p-STAT3 (Y705) and PD-L1 in HepG2215 cells after treatment with NSC-74859 for 24 h. JAK1, STAT3 and p-STAT3 (Y705) in sh*YAP1*-HepG2215 cells and the liver tumors in *Yap1*^LKO^ mice. \**P* < 0.05. D. YAP1 promoted PD-L1 expression by JAK1/STAT1,3 pathways. E. Flow cytometry results of the percentage of PD-1^−^CD8^+^ and PD-1^+^CD8^+^ T cells in the spleen in verteporfin-treated C57BL/6 mice. F. IF results of CD8^+^ T and CD4^+^ T cells in the liver tumor tissues of *Yap1*^LKO^ mice. \**P* <0.05 *vs* DMSO group. G. TGF-β level in liver tissues in verteporfin-treated C57BL/6 mice by ELISA.

Further, *YAP1* knockdown decreased PD-L1 expression in HepG2215 cells (Fig.5A), suggesting that YAP1 promoted PD-L1 expression. JAK1-STAT1 signing is the main pathway responsible for IFN-γ induced PD-L1 expression in HCC cells (Li et al. 2018). Meanwhile, YAP1 interacted with TEA domain transcription factor (TEAD) to regulate JAK-STAT pathway (Gruber et al. 2016). We found that *Yap1* knockout restricted the expressions of STAT1, p-STAT1 (T727) and p-STAT3 (Y705) in liver tumors of *Yap1*^LKO^ mice (Fig. 5B, and 5C). But, the expression level of STAT3 did not change in *Yap1*^LKO^ mice and sh*YAP1*-HepG2215 cells (Fig. 5C). These results showed that YAP1 promotes p-STAT1 (T727) and p-STAT3 (Y705) activation, not STAT3 expression. JAK1, but not JAK2, is the primary mediator of JAK-STAT pathway in melanoma or bladder tumor (Luo et al. 2018, Daza-Cajigal et al. 2019). Interestingly, JAK1, the upstream molecule of JAK-STAT pathway was also reduced in *Yap1*^LKO^ mice (Fig. 5B and 5C). These results showed that YAP1 promotes JAK1 expression.

p-STAT3 (Y705) upregulated the expression level of PD-L1 (Bu et al. 2017). Further, the expression of PD-L1 decreased after treated with p-STAT3 (Y705) inhibitor, NSC74859 (Fig. 5D). YAP1 knockout inhibited the expression of p-STAT3 (Y705) in liver tumors of *Yap1*^LKO^ mice (Fig. 5C). These results showed that YAP1 interacted with p-STAT3 (Y705) to promoted PD-L1 expression. Taken together, we suggested that JAK1/STAT1, 3 facilitate PD-L1 expression depending on YAP1 (Fig. 5E).

Besides PD-L1 on the tumor cells, PD-1^+^CD8^+^ T cells were correlated with exhausted signature in HCC (Ma, Zheng, et al. 2019). YAP1 inhibitor, verteporfin, decreased the percentage of PD-1^+^CD8^+^ T cells, while increased the percentage of PD-1^−^CD8^+^ T cells in spleen in DEN/TCPOBOP-induced liver tumor mice (Fig. 5F). The result suggested that YAP1 increased the number of exhausted CD8^+^ T cells in spleen. Further, we detected that the number of CD4^+^ T and CD8^+^ T cells were elevated in liver tumor of *Yap1*^LKO^ mice (Fig. 5B and 5G). The result suggested that YAP1 reduced the number of T cells in liver tumor niche. TGF-β inhibited CD8^+^ T cell activation and promoted Treg differentiation (Ringelhan et al. 2018). Verteporfin decreased the expression level of TGF-β in liver tumor of C57BL/6 mice (Fig. 5H). These results showed that YAP1 knockout alleviated suppressive tumor microenvironment *in vivo*.

### DHA broke the tumor immunosuppressive microenvironment

Further, anti-PD-1 treatment increased the percentage of PD-1^+^CD4^+^ and PD-1^+^CD8^+^ T cells, decreased PD-1^−^CD8^+^ T cells in PBMC of C57BL/6 mice with DEN/TCPOBOP-induced liver tumor (Fig. 6A). And anti-PD-1 increased the percentage of PD-1^+^ CD4^+^ T cells, decreased PD-1^−^CD4^+^ and PD-1^−^CD8^+^ T cells in spleen (Fig. 6B). These results suggested that anti-PD-1 treatment increased the number of exhausted T cells and decreased the functional T cells in peripheral blood and spleen. Furthermore, anti-PD-1 treatment decreased IFN-γ in liver tumor tissues (Fig. 6D).

**Figure 6.**
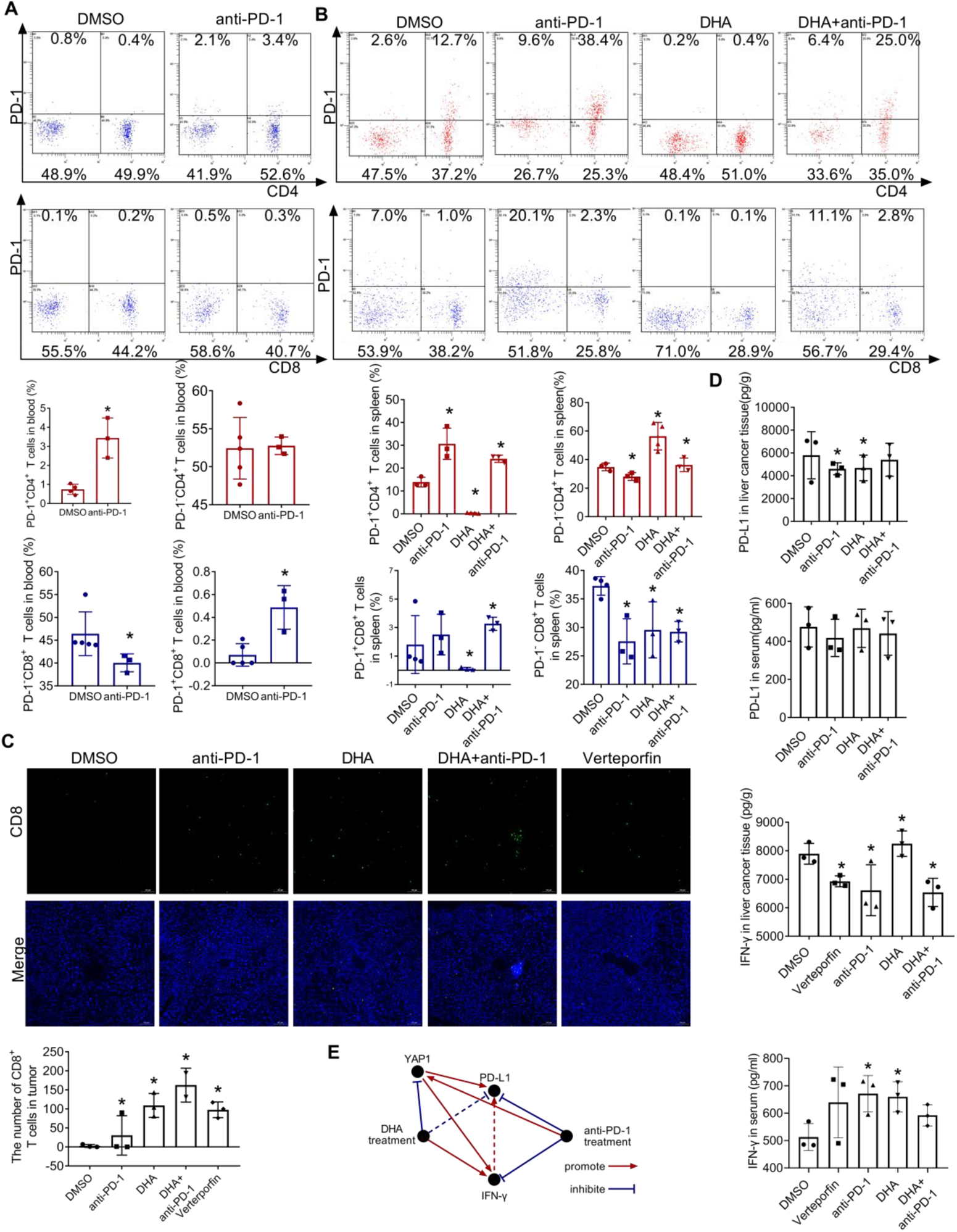
DHA broke the tumor immunosuppressive niche. A. The percentage of exhausted T cells (PD-1^+^CD4^+^and PD-1^+^CD8^+^) in PBMC from anti-PD-1-treated C57BL/6 mice by flow cytometry. \**P* <0.05 *vs* DMSO group. B. Flow cytometry result of the percentage of PD-1^−^ CD4^+^, PD-1^+^CD4^+^, PD-1^−^ CD8^+^ and PD-1^+^ CD8^+^ T cells in spleen from C57BL/6 mice after anti-PD-1 and/or DHA treatment. \**P* <0.05 *vs* DMSO group. C. IF results of CD8^+^ T cells in the liver tumor tissues in anti-PD-1 and/or DHA or verteporfin treated mice. \**P* <0.05 *vs* DMSO group. D. IFN-γ and PD-L1 in liver tumor tissues or in the serum by ELISA. \**P*<0.05 *vs* DMSO group. E. Schematic model of the regulatory pathway and mechanism of YAP1 and DHA in tumor immune evasion during anti-PD-1 therapy.

Interestingly, DHA decreased the percentage of PD-1^+^CD4^+^ T cells and PD-1^+^ CD8^+^ T cells, while increased the percentage of PD-1^−^CD4^+^ T cells in spleen (Fig. 6B). And DHA decreased PD-L1 in liver tumor tissues and increased IFN-γ in serum and liver tumor tissues (Fig. 6D).

Furthermore, DHA increased CD8^+^ T cells in liver tumor tissues (Fig. 6C), similarly to the result of *Yap1* ^LKO^ (Fig. 5G) or verteporfin-treated mice (Fig. 6C). These results suggested that DHA inhibited YAP1, leading to improvement of the anti-tumor immune microenvironment in mice.

Notably, DHA combined with anti-PD-1 decreased the percentage of PD-1^+^ CD4^+^ T cells and increased the percentage of PD-1^−^CD4^+^ T cells compared with anti-PD-1 treatment (Fig. 6B). However, the combination treatment was not significantly changed the percentage PD-1^+^CD8^+^ and PD-1^−^CD8^+^ T cells compared with anti-PD-1 treatment (Fig. 6B). Furthermore, the number of CD8^+^ T cells of tumor tissues were increased in DHA combined with anti-PD-1 treatment compare with DHA, anti-PD-1 or verteporfin alone (Fig. 6C). Together, these data demonstrated that DHA combined with anti-PD-1 treatment promoted CD4^+^ T cell activation in the spleen and increased the number of CD8^+^ T cells in tumor tissues.

## Discussion

Single-agent anti-PD-1 therapy was far from enough to improve the survival rate of HCC patients. Here, we present evidenced that YAP1 in tumor tissues directly promotes PD-L1 and reduced CD4^+^ and CD8^+^ T cells in the local tumor tissues. Indirectly, JAK1/STAT1, 3 promoted PD-L1 expression depending YAP1. Notably, we suggested that DHA, a drug approved by FDA, acted as a YAP1 inhibitor, broke the immunosuppressive microenvironment, leading to increase the efficacy of anti-PD-1 therapy in mice.

JAK-STAT pathway also causes ICI resistance in some melanoma patients (Nguyen et al. 2021). p-STAT3 (Y705) was detected in approximately 60% of HCC samples (He et al. 2010). STAT3 directly binds to the PD-L1 promoter to increase its expression (Marzec et al. 2008). Interestingly, we showed that p-STAT3 (Y705) was reduced in HepG2215 cells and liver tumor cells of *Yap1^LKO^* mice. YAP1 still binds to STAT3 promoter in nucleus and promotes STAT3 expression at the transcriptional level in pancreatic ductal adenocarcinoma cells (Gruber et al. 2016). STAT1 is also highly expressed in HCC tissues (Ma, Chen, et al. 2019). Individual STAT1 and STAT3 activation induces PD-L1 expression in HCC cells (Garcia-Diaz et al. 2017). In addition, p-STAT1 (T727) and p-STAT3 (Y705) dimerized and bound to PD-L1 promoter, leading to PD-L1 expression in human breast cancer cells (Sasidharan Nair et al. 2018). Our results suggest that p-STAT1 (T727) and p-STAT3 (Y705) promoted PD-L1 expression by YAP1 in liver tumor cells, separately. Furthermore, JAK1, but not JAK2, is the primary mediator of JAK/STAT pathway associated with PD-L1-mediated immune surveillance in melanoma (Luo et al. 2018) and bladder cancer (Daza-Cajigal et al. 2019). Accordingly, IL-6-activated JAK1 phosphorylates PD-L1 and induces PD-L1 glycosylation, which maintain PD-L1 stability in liver tumor cells (Chan et al. 2019). STAT3 activation in hepatocytes is not required for the tumor formation after knockout of 2 mammalian Hippo kinases *(Mst1* and *Mst2)*, which inhibit YAP1 activation in mice (Kim et al. 2017). Moreover, our results suggested that YAP1 knockout restrains JAK1 expression in *Yap1^LKO^* mice. HCC is a cancer with high percentage (~7%) of JAK1 mutations (Kan et al. 2013). Especially, simply knockdown YAP1 can induce PD-L1 expression in HCC.

YAP1 directly bound to PD-L1 promoter (Kim et al. 2018). YAP1 knockdown decreased PD-L1 expression in HepG2215 cells. Consistently, our previous research also found that verteporfin inhibited PD-L1 expression in liver tumor cells (Li et al. 2020). However, anti-PD-1 treatment increased YAP1 and decreased PD-L1 in liver tumor in mice, suggesting others besides YAP1 can be involved in PD-L1 expression. IFN-γ was an important cause of inducing PD-L1 expression in tumor microenvironment (Qian et al. 2018, Thiem et al. 2019). Here we showed that anti-PD-1 treatment decreased PD-L1 and IFN-γ expression in liver tumor tissues of mice. In addition, interactions between increased YAP1 and TGF-β in hepatocytes stimulate epithelial-to-mesenchymal transition (EMT)-like response in a TGF-β-enriched microenvironment after partial hepatectomy (Oh et al. 2018). Further, verteporfin suppressed the TGF-β expression in the liver tumor tissue of mice. Therefore, elevated YAP1 is involved in HCC tumor microenvironment during anti-PD-1 treatment.

Notably, selectively knockout *Yap1* from hepatocytes increased CD4^+^ and CD8^+^ T cell infiltration in liver tumor niche of *Yap1^LKO^* mice. Verteporfin, a YAP1 inhibitor to disrupt the interaction between YAP/TAZ and TEAD complex, increased the percentage of CD8^+^ T cells in liver tumor niche in mice. Consistently, verteporfin treatment increased T cell activation without significant effect on T cell proliferation (Stampouloglou et al. 2020). Collectively, disruption of YAP1 in liver tumor cells recruits CD8^+^ T cells to tumor niche. Meanwhile, YAP1 in T cells is elevated upon T-cell activation, and deletion of YAP1 in T cells promotes T-cell infiltration into the local tumor niche (Stampouloglou et al. 2020). Therefore, we considered that YAP1 inhibitor reduced PD-L1 expression on tumor and increase T cell recruitment.

DHA was a derivative of artemisinin extracted from *artemisia annua Linn* (Guo 2016). Our previous studies showed that DHA inhibited HepG2(Hao et al. 2021) and HepG2215 cells (Shi et al. 2019). Meanwhile, some studies suggested that DHA downregulated PD-L1 expression in non-small cell lung cancer (Zhang et al. 2020). Interestingly, DHA decreased YAP1 in tumor tissues but increased YAP1 in para-tumor, leading to the increased ratio of YAP1, suggesting the competitive advantage of para-tumor tissues, which appears to eliminate liver tumor cells in mice (Moya et al. 2019). The result is similar to YAP1 inhibitor, verteporfin. Further, we showed that the ratio of YAP1 expression was significantly negatively correlated with YAP1 in tumor tissues from HCC patients. Similarly, PD-L1 and CD8 were decreased in tumor tissues compared to adjacent normal liver tissues from 143 HCC patients (Guo et al. 2020). We further showed that combination anti-PD-1 with DHA increased in para-tumor and decreased YAP1 in tumor tissues. However, anti-PD-1 treatment increased YAP1 expression in para-tumor and tumor tissues, and the ratio of YAP1. Confusingly, anti-PD-1 treatment increased the number of exhausted T cells and decreased the functional T cells in blood and spleen of HCC mice. It was generally accepted that successful anti-tumor immune responses following anti-PD-1 therapy required tumor-specific CTLs in the tumor niche (Wu et al. 2019). Interestingly, DHA increased the number of CD8^+^ T cells in tumor tissues, about 3-fold change that of anti-PD-1 treatment. This effect is similar to that of verteporfin treatment and *Yap1*^LKO^ mice. In addition, DHA decreased the number of exhausted

T cells (PD-1^+^CD4^+^ or PD-1^+^CD8^+^ T cells), increased the functional PD-1^−^CD4^+^ cells in spleen.

Consistently, DHA induced IFN-γ^+^CD8^+^ T cell proliferation, and reduced the number of CD4^+^CD25^+^Foxp3^+^ T cells in melanoma tumor tissue compared with normal tissue (Yu et al. 2020). Moreover, the advantages of combination therapy over treatment alone were reduced YAP1 in tumor tissue and increased the ratio of YAP1 in adjacent tissues to inhibit the tumor volume. Notably, the combination of DHA and anti-PD-1 was increased CD8^+^ T cell number in tumor tissues. Therefore, DHA was used as a potent YAP1 inhibitor and combined with anti-PD-1 to suppression of immune evasion in liver tumors.

In summary, we confirmed firstly that YAP1 knockdown in liver tumor cells suppressed PD-L1 expression and recruit CTLs, leading to break immune evasion in tumor niche. Mechanistically, JAK1/STAT1, 3 promoted PD-L1 expression by YAP1 in HCC. Finally, DHA, as a potent YAP1 inhibitor, broke the immunosuppressive niche in liver tumor tissues to improve the effect of anti-PD-1 therapy.

## Materials and Methods

### Bioinformatics analysis

Raw counts of RNA-sequencing data, corresponding clinical information from 371 HCC and normal tissue samples were obtained from The Cancer Genome Atlas (TCGA) (portal.gdc.cancer.gov/) (Weinstein et al. 2013). The mRNA expression level of *YAP1* was analyzed compared with normal samples. And the expression of *YAP1* in different tumor stages of HCC was analyzed.

### Cell culture

HepG2215 cells were purchased from American Type Culture Collection (Manassas, VA, USA). sh*YAP1*-HepG2215 cells has been constructed previously (Li et al. 2020). They were cultured in DMEM (Gibco/Thermo Fisher Scientific, Beijing, China) supplemented with 10% fetal bovine serum (Gibco/Thermo Fisher Scientific, Beijing, China), 100 U/ml penicillin and 100 ug/ml streptomycin at 37 °C and 5% CO_2_, in an atmosphere of 100% humidity.

### Cell treatment

NSC-74859 (MCE, China) was dissolved in DMSO (Sigma-Aldrich, USA) and stored at −20°C. HepG2215 cells were treated with NSC-74859 (100μM) for 24 h. The culture medium containing DMSO was used as the control.

### Animal experiments

The protocol was approved by the Ethics Committee for Animal Experiment of Hebei University of Chinese Medicine (Shijiazhuang, China) (Permit number: YXLL2018002). All animal were maintained in the SPF facility with constant temperature (22-24 °C) and a dark-light cycle of 12h/12h. All experiments were conducted with male mice. C57BL/6 mice (Vital River Laboratory Animal Technology Co. Ltd., Beijing, China) at the age of 3 weeks were used. The genetically engineered albumin-cre mice were purchased from Guangzhou Cyagen Biosciences (Guangzhou, China). *Yap1*^flox/flox^ mice with a *loxP*-flanked *Yap1* allele on a C57BL/6N background were generated. Albumin-cre mice were crossed with *Yap1*^flox/flox^ mice to produce *Yap1*^flox/flox, Alb-cre^ (mark as *Yap1*^LKO^) mice, the liver-specific knockout *Yap1* mice. DNA was extracted from mice tail, and amplified PCR in a thermocycler machine for genotype identification. The primer sequences are shown in supplementary Table 1.

### DEN/TCPOBOP-induced liver tumor model in C57BL/6 mice

Each 3-week-old male C57BL/6 mice, including wild type, *Yap1*^flox/flox^ and *Yap1*^LKO^ mice, was injected intraperitoneally with N-nitrosodiethylamine (DEN) at the dose of 25 mg/kg body weight.

At the age of 4 weeks, the mice received ten consecutive biweekly injections with TCPOBOP (3 mg/kg). The method was introduced as previously described (Li et al. 2020, Bergmann et al. 2017). At the 24th week of age, the tumors in liver were determined by ultrasound in the mice by an imaging system (Vevo 2100, VisualSonics Inc., Toronto, Canada) with an MS250 ultrasound transducer (Li et al. 2020). After successful modeling, they were randomly divided into five groups (n=6) and treated for 25 d. During the treatment, no mice died from tumor loading. The mice of anti-PD-1 group were injected intraperitoneally with anti-PD-1 (BioXcell, USA, 10 mg/kg) every 3 d. DHA group were intraperitoneally injected daily with DHA in DMSO (25mg/kg) (Shi et al. 2017). The mice in anti-PD-1 combined with DHA group were intraperitoneally injected with DHA (25mg/kg) and anti-PD-1 (10 mg/kg) once every 3 d. Verteporfin group were intraperitoneally injected daily with verteporfin in DMSO (100 mg/kg) for 25 days. The mice in the control group were intraperitoneally injected daily with 0.1% DMSO in 100μl physiological saline.

### Quantitative reverse transcription-polymerase chain reaction (qRT-PCR)

Total RNA was extracted from *shYAP1-*HepG2215 cells using TRIzol reagent (ThermoFisher Scientific, America) according to the manufacturer’s instructions. Then, RNA was converted into cDNA with Prime Script RT reagent kit (Takara Bio Inc, Japan). Real time fluorescence qRT-PCR was performed on a Real-Time PCR system (BIOER Co. Ltd., Kokyo, Japan). Finally, the Ct values were obtained from the ABI 7500 fast v2.0.1 software. The ΔΔCt method was used to represent mRNA fold change. The primer sequences are shown in supplementary Table 1.

### Western blot

Total protein was harvested from HCC cells using column tissue and cell protein extraction kit (Shanghai Yamei Biotechnology Co., Ltd, Shanghai, China). Proteins were separated on 12% SDS-PAGE and transferred to PVDF membranes. After blocking, PVDF membranes were incubated with primary antibodies, and then secondary antibodies. The primary antibodies were rabbit anti-YAP1 antibody (CST, #14074), rabbit anti-TAZ antibody (CST, #72804), rabbit anti-JAK1 antibody (Abways, CY7173), mouse anti-STAT3 antibody (CST, #9139), mouse anti-p-STAT3(Y705) antibody (CST, #4113), rabbit anti-STAT1 antibody (CST, #14994), mouse anti-PD-L1 antibody (abcam, ab238697), rabbit anti-GAPDH antibody (Abways, ab0037), rabbit ACTIN antibody (Abways, ab0035), and rabbit anti-Tubulin antibody (Abways, ab0039). The secondary antibodies were goat anti-rabbit IgG-HRP (ZSGB-BIO, ZB-2301) and goat anti-mouse IgG-HRP (Abways, ab0102). The bands were visualized by enhanced chemiluminescence (ECL) kit (Shanghai Share-bio Technology Co., Ltd, China) and detected by Fusion FX5 Spectra ECL detection systems (Vilber, France). The band intensity was measured by the Image-Pro Plus v6.0 software (Media Cybernetics. USA).

### Immunohistochemistry (IHC)

Human HCC tissue microarray was purchased from Servicebio (China, Wuhan, no: IWLT-N-64LV41 Live C-1401). The liver tissue from the mice was fixed with 4% paraformaldehyde and embedded in paraffin. Then, the liver tissue sections (2 μm in thickness) were deparaffinised, and dehydrated before staining with haematoxylin and eosin (H&E). After deparaffinized and rehydrated, tissue sections were retrieved antigen, inactivated the endogenous enzyme, and incubated with primary antibodies (rabbit anti-p-STAT1 (T727) (abcam, ab109461), rabbit anti-Ki67 (CST, 12202S) and other antibodies have been described in the Western blot section). PBS was used as the negative control for the primary antibody. And then, the sections were rinsed and incubated with the secondary antibody. Subsequently, the sections were developed with 3, 3-diaminobenzidine (DAB) kit (ZSGB-Bio, China). Finally, the cytomembrane/cytoplasm stained with light yellow or tan were regarded as positive cells.

IHC staining was scored according to the following method. The percentage of positive cells in total cells of ≤ 5% was Negative expression (-) and scored as 0 point. 6-25% was weak expression (+) and 1 point. 26-50% was moderate expression (++) and 2 points. > 50% was strong expression (+++) and 3 points. The judgment of protein expression is based on both the staining intensity and positive cell rate, and the product of these two values was calculated. After the multiplication of the two scores, they were divided into two groups: the group with the product of not less than 3 points was defined as the high expression group, and the group with the product of less than 3 points was defined as the low expression group.

### Immunofluorescence assay (IF)

After deparaffinized and rehydrated, tissue sections from mice were retrieved antigen, inactivated the endogenous enzyme, incubated with primary antibody, and then second antibody. The primary antibodies were rabbit anti-CD4 (Seville Biological, GB11064,) and rabbit anti-CD8 (Seville Biological, GB13429). The secondary antibody was FITC goat anti-rabbit IgG-HRP (Seville Biological, GB22303,). Finally, the tissue sections were washed and incubated in DAPI. All fluorescence images were observed under a Biosystem microscopy (Leica, Wetzlar, Germany).

### Flow cytometry

The mouse blood and single cells from spleen were used to separate Peripheral blood mononuclear cells (PBMC) with Mouse peripheral blood lymphocyte isolation solution kit (P8620, Solarbio, Beijing, China). The following fluorochrome-labeled mono-antibodies and staining reagents were used according to the protocols. Cells from PBMC or spleen were stained with anti-mouse CD3ɛ, FITC (MultiSciences, AM003E01), anti-mouse CD279 (PD-1), APC (Biolegend, 135210), anti-mouse CD8a, PE (MultiSciences, AM008A04), and anti-mouse CD4, PE (MultiSciences, AM00404). The cells were analyzed with by an FC 500 MCL flow cytometer (Beckman Coulter, Inc. USA) and analyzed CXP software (version 2.1; Beckman Coulter, Inc.).

### Enzyme linked immunosorbent assay (ELISA)

Tissue homogenates of liver tumor or serum were added to 96-well plates from these kits, including Mouse PD-L1 ELISA Kit (mlbio, ml058347), Mouse TGF-β ELISA Kit (mlbio, ml057830), Mouse Alanine Aminotransferase (ALT) ELISA Kit (mlbio, ml063179), Mouse IFN-γ ELISA Kit (mlbio, ml002277). The main steps are carried out according to the instructions. Finally, the Stop Solution changes the color from blue to yellow, and the intensity of the color is measured at 450 nm using a spectrophotometer (Rayto, Shenzhen, RT-6100). The concentration of the target protein in the samples is then determined by comparing the O.D. of the samples to the standard curve.

### Statistical analysis

Statistical analyses were performed by SPSS 23.0 statistics software (SPSS, Chicago, IL) and GraphPad Prism 8 software. All *in vitro* and *in vivo* experiments were repeated at least three times and at least three samples were taken at a time. If data were normally distributed, they are represented as means ± SD. When more than two groups were included, one-way analysis of variance would be used. Differences were considered statistically significant when *P*-value was less than 0.05.

## Acknowledgements

This work was financially supported by the National Natural Science Foundation of China (81873112).

## Author contributions

Xinli Shi designed research. Qing Peng, Shenghao Li, Yinglin Guo, Liyuan Hao, Zhiqin Zhang, Jingmin Ji, Yanmeng Zhao, Caige Li, Yu Xue and Yiwei Liu performed research. Qing Peng, Shenghao Li, Yinglin Guo, and Liyuan Hao analyzed data. Xinli Shi, Qing Peng and Shenghao Li wrote the paper.

## Figure legends

**Figure S1.**
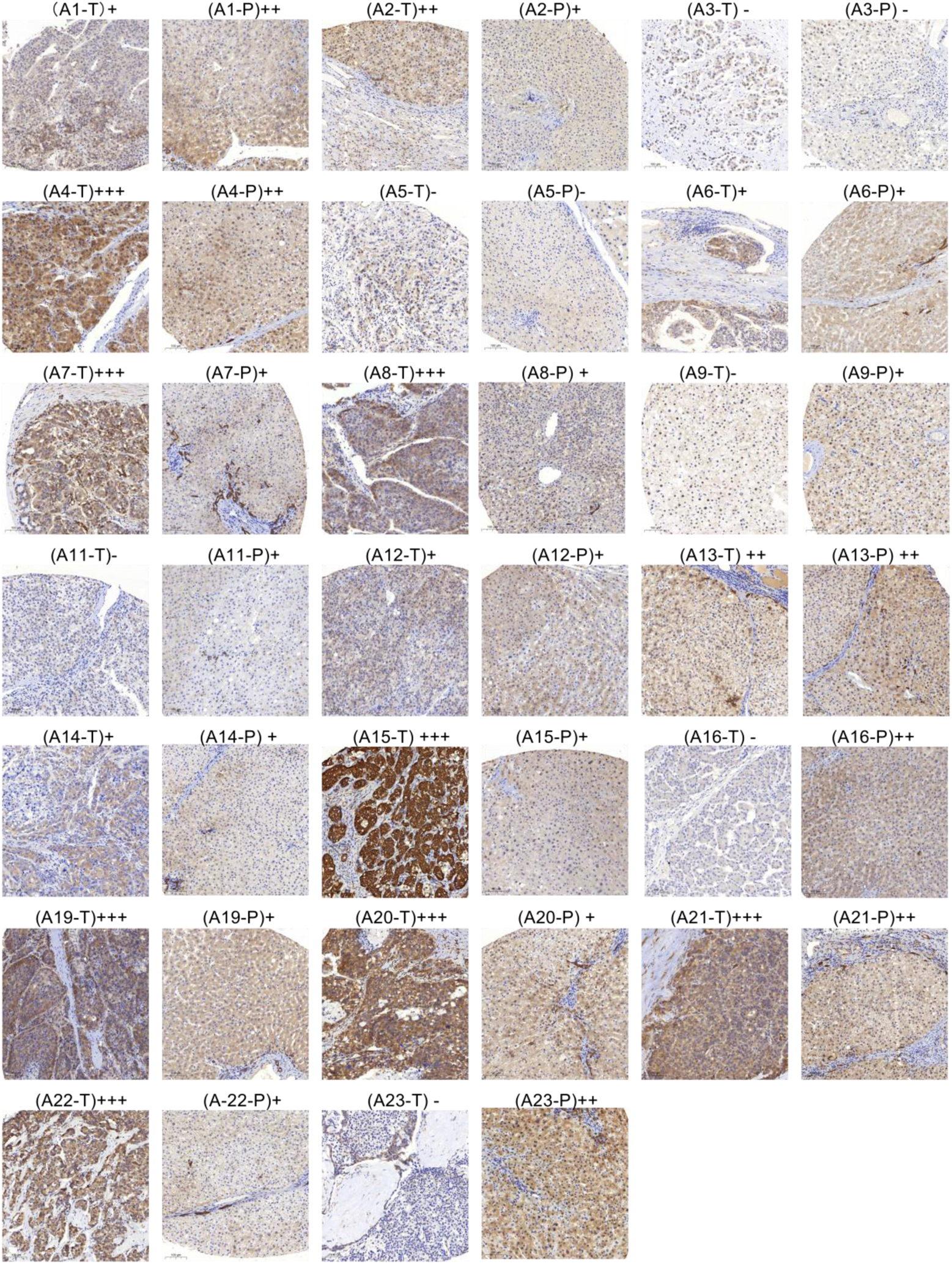
The expression of YAP1 in tumor (T) and para-tumor tissues (P) by the HCC tissue microarray. Axx indicate patient number.

**Figure S2.**
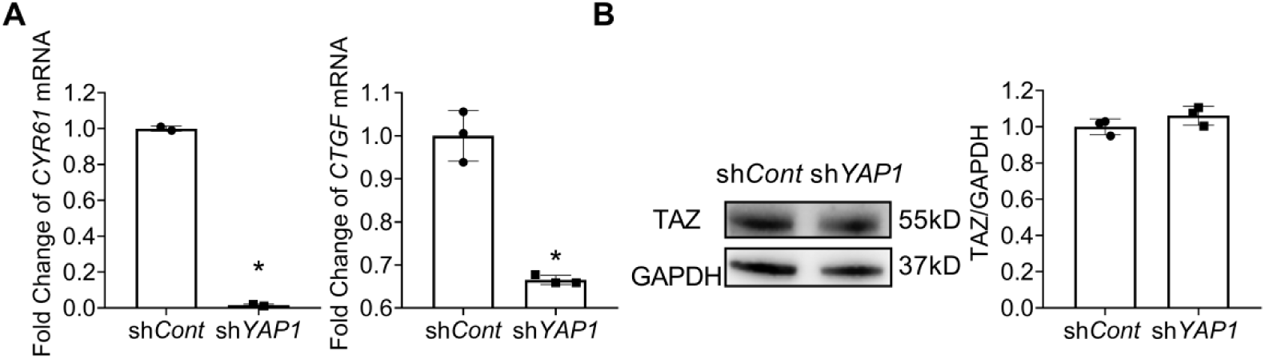
YAP1 and the downstream genes CYR61 and CTGF were decreased, while TAZ did not change in shYAP1-HepG2215 cells. A. The relative mRNA levels of *CYR61*and *CTGF* in shYAP1-HepG2215 cells by q-PCR. \**P* <0.05 vs DMSO group. B. Western blot results of TAZ of the liver tumor in sh*YAP1*-HepG2215 cells.

**Supplementary Table 1.**
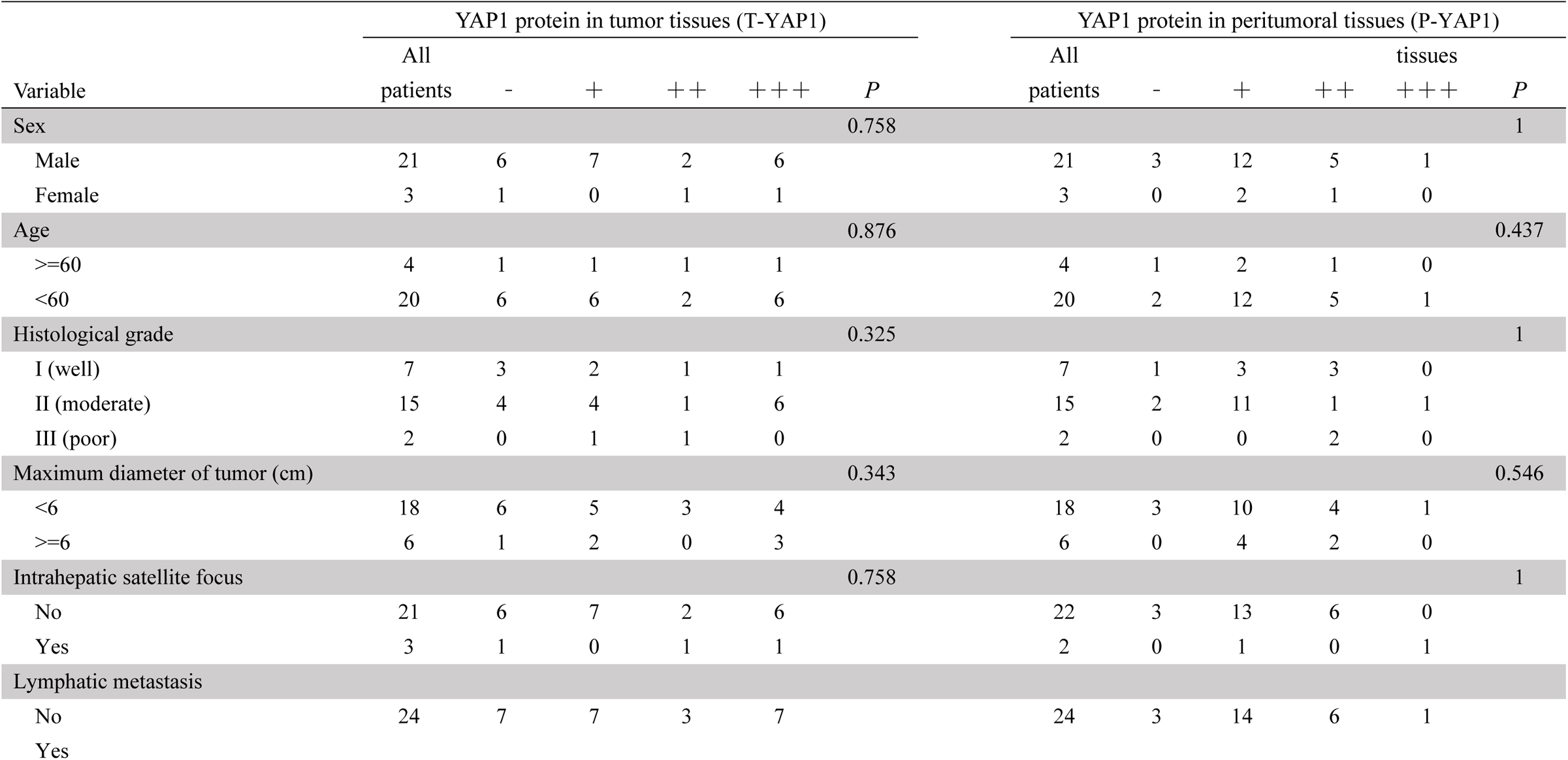
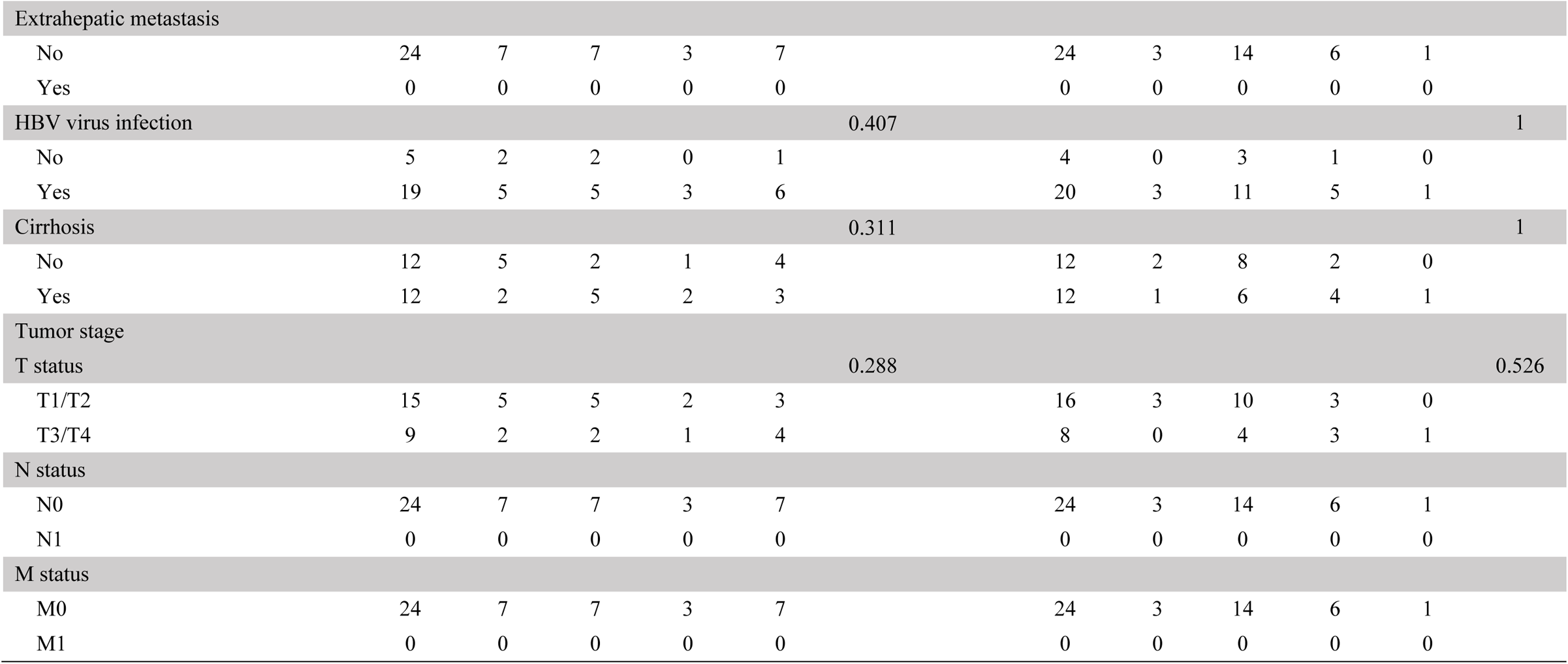
Association of YAP1 expression with the clinicopathological factors of patients with HCC in tumor and peritumoral tissues expression in Human HCC tumor tissue microarray.

**Supplementary Table 2.**
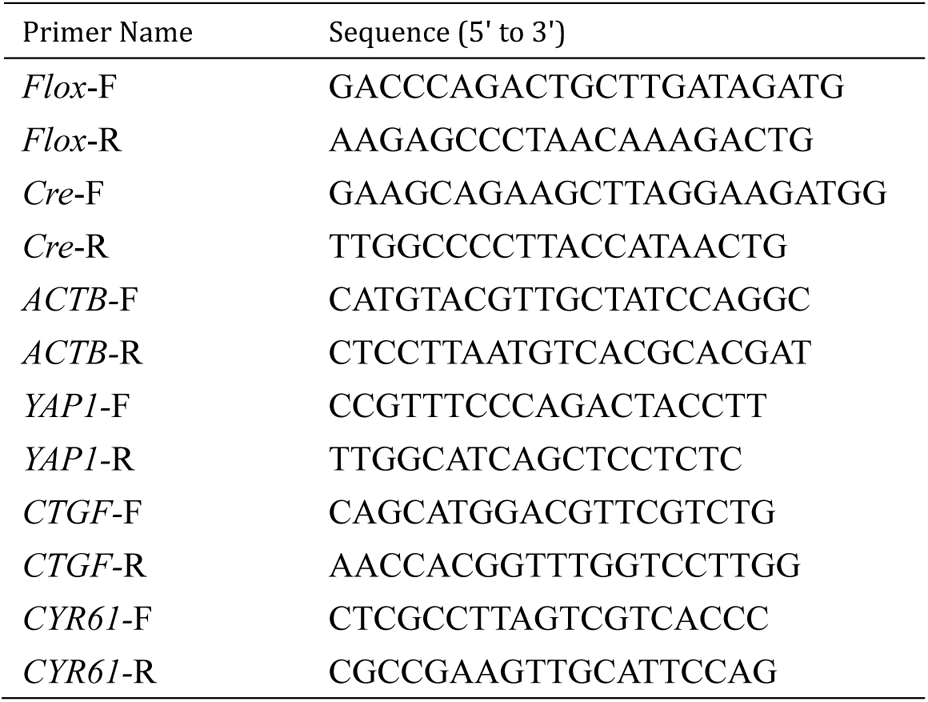
Primers for cloning

